# Dynamic hormone control of stress and fertility

**DOI:** 10.1101/2020.08.24.264234

**Authors:** Eder Zavala, Margaritis Voliotis, Tanja Zerenner, Joël Tabak, Jamie J Walker, Xiao Feng Li, John R Terry, Stafford L Lightman, Kevin O’Byrne, Krasimira Tsaneva-Atanasova

## Abstract

Neuroendocrine axes display a remarkable diversity of dynamic signalling processes relaying information between the brain, endocrine glands, and peripheral target tissues. These dynamic processes include oscillations, elastic responses to perturbations, and plastic long term changes observed from the cellular to the systems level. While small transient dynamic changes can be considered physiological, larger and longer disruptions are common in pathological scenarios involving more than one neuroendocrine axes, suggesting that a robust control of hormone dynamics would require the coordination of multiple neuroendocrine clocks. The idea of apparently different axes being in fact exquisitely intertwined through neuroendocrine signals can be investigated in the regulation of stress and fertility. The stress response and the reproductive cycle are controlled by the Hypothalamic-Pituitary-Adrenal (HPA) axis and the Hypothalamic-Pituitary-Gonadal (HPG) axis, respectively. Despite the evidence surrounding the effects of stress on fertility, as well as of the reproductive cycle on stress hormone dynamics, there is a limited understanding on how perturbations in one neuroendocrine axis propagate to the other. We hypothesize that the links between stress and fertility can be better understood by considering the HPA and HPG axes as coupled systems. In this manuscript, we investigate neuroendocrine rhythms associated to the stress response and reproduction by mathematically modelling the HPA and HPG axes as a network of interlocked oscillators. We postulate a network architecture based on physiological data and use the model to predict responses to stress perturbations under different hormonal contexts: normal physiological, gonadectomy, hormone replacement with estradiol or corticosterone (CORT), and high excess CORT (hiCORT) similar to hypercortisolism in humans. We validate our model predictions against experiments in rodents, and show how the dynamic responses of these endocrine axes are consistent with our postulated network architecture. Importantly, our model also predicts the conditions that ensure robustness of fertility to stress perturbations, and how chronodisruptions in glucocorticoid hormones can affect the reproductive axis’ ability to withstand stress. This insight is key to understand how chronodisruption leads to disease, and to design interventions to restore normal rhythmicity and health.

## 1 INTRODUCTION

A robust dynamic interplay between body rhythms is essential to sustain healthy states. This requires the coordination of several regulatory systems spanning multiple levels of organisation, from molecular, to cellular, to the whole organism. Neuroendocrine axes are the perfect example of such interlocked-regulatory systems controlling body rhythms, with the brain decoding circadian and stress inputs as well as integrating feedback signals from endocrine organs. The hypothalamic-pituitary-adrenal (HPA) axis and the hypothalamic-pituitary-gonadal (HPG) axis are the major neuroendocrine systems underpinning stress and fertility, respectively. These axes control a range of hormonal and neural activity rhythms exhibiting ultradian (*<*24h), circadian (∼24h) and infradian (>24h) periodicity [1], as well as responses to environmental, biological and behavioural perturbations. For example, the HPA axis uses feedback loops to regulate stress responses while sustaining ultradian and circadian glucocorticoid (CORT) rhythms [2, 3]. On the other hand, the HPG axis controls infradian oscillations of reproductive hormones secreted in response to changes in the ultradian frequency of gonadotropin-releasing hormone (GnRH). GnRH secretion is controlled by a hypothalamic pulse generator (PG) [4], which is in turn modulated by reproductive hormones (Fig. 1A). Mathematical modelling has significantly contributed to our understanding of the origin of this rhythmic behaviour [3, 4, 5, 6, 7], as well as the ability of these systems to respond to perturbations.

**Figure 1.**
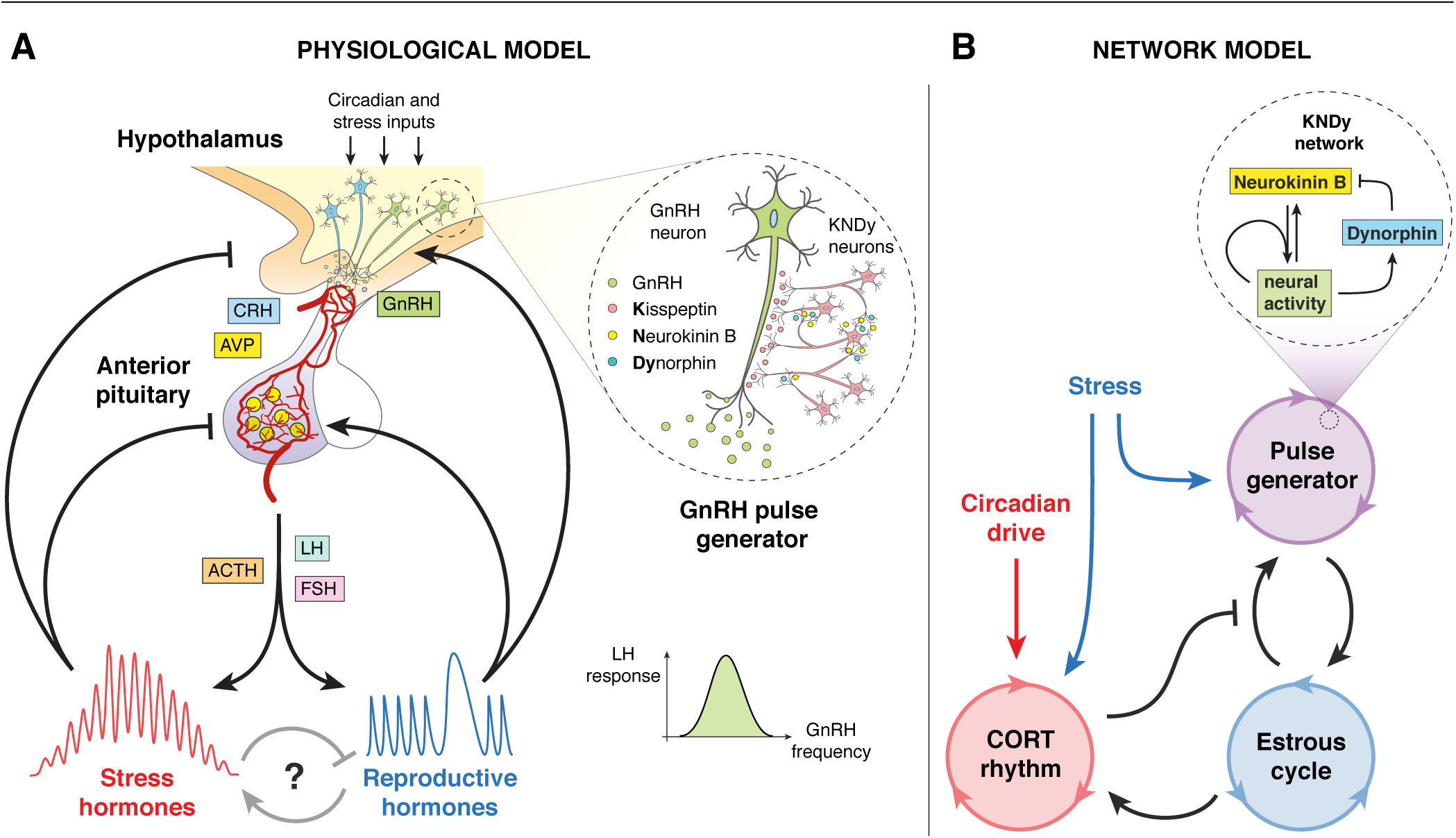
Pictorial representation of the model. **(A)** Physiological model of the stress and reproductive neuroendocrine axes controlling ultradian, circadian and infradian hormone oscillations. Includes the KNDy neuronal network controlling the GnRH pulse generator. Adapted from [5]. **(B)** Network model of the systems-level cross-regulation between glucocorticoid (CORT) rhythms, the hypothalamic GnRH pulse generator and the estrous cycle, subject to stress and circadian inputs. Includes a mean-field model of the KNDy network from [4].

In recent years, animal models have greatly enhanced our understanding of how specific perturbations in the HPA axis dynamics affect the HPG axis [8, 9, 10, 11], as well as how the activity of the HPG axis can in turn modulate the dynamics of the HPA axis [12, 13, 14, 15]. Furthermore, human studies have shown how glucocorticoid excess can have profound effects on the menstrual cycle [16, 17, 18]. Many studies have shown the links between stress and fertility [19, 20, 21, 22] and provided evidence of cross-regulatory interactions between the HPA and HPG axes [23, 24]. However, there is still a limited understanding of whether and how the HPA and HPG axes coordinate their hormone rhythms, how perturbations to one axis impact upon the other, what makes their dynamics robust to such perturbations, and in what circumstances chrono-disruptions can lead to disease.

In this manuscript, we investigate the dynamic control of stress and fertility by means of a mathematical model that accounts for the complex interactions between the HPA and HPG axes. First, we postulate that these neuroendocrine systems behave as a network of coupled oscillators that coordinate ultradian, circadian and infradian rhythms, and validate the model predictions against physiological observations in female rodents. Second, we consider the evidence on stress-induced suppression of GnRH pulse generator activity dependent of estradiol, and use the model to understand the role of estradiol-dependent effects on the HPA axis [12, 25]. We also explore the simultaneous effect of exogenous estradiol and glucocorticoids on the dynamics of the GnRH pulse generator [8]. Third, we use the model to explore how perturbations such as stressors and chronic changes in gonadal steroids and glucocorticoid levels can disrupt normal rhythmicity and lead to dysregulations that propagate from one neuroendocrine system to the other. To do so, we consider typical restraint stress signals [26] to brain regions that are connected to the hypothalamus, thus affecting both the HPA and HPG axes [20, 27]. Importantly, our model considers the signalling role of regulatory neuropeptides (e.g., Neurokinin-B and Dynorphin) within the KNDy neural network in stress-induced suppression of the GnRH pulse generator [4, 28, 29], which has implications for our understanding of how stress signals are decoded by the reproductive axis. Lastly, we predict an increase of the estrous cycle length under hiCORT and discuss how our model can help understand the mechanisms allowing robust control of ovulation despite the effect of stressors.

## 2 MODEL AND METHODS

### 2.1 Mathematical modelling

We focus on the systems-level outputs and cross-regulation of the stress and reproductive axes, which in turn we model as a network of coupled oscillators (Fig. 1B). We modelled this through a system of Ordinary Differential Equations (ODEs), where each oscillator represents an aspect of neuroendocrine rhythmic activity that can be characterised by a phase *φ*, a frequency *ω*, and an amplitude *A*. Our model consists of a master circadian oscillator in the hypothalamus, a glucocorticoid (CORT) oscillator with ultradian rhythmicity driven by the circadian oscillator, a pulse generator oscillator governed by the Kisspeptin, Neurokinin B, Dynorphin (KNDy) network regulating pulses of GnRH secretion, and an oscillator representing the estrous cycle. The equations for these oscillators are listed below, with coupling functions, parameter values, and further details of the model development described in the Supplementary Material.

#### Circadian cycle

A fixed period hypothalamic oscillator to control the circadian rhythm of CORT:

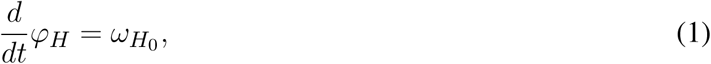

where *φ*_*H*_ is the hypothalamic phase and 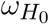 is the natural frequency of the hypothalamic circadian drive.

#### CORT oscillator

Accounts for CORT ultradian oscillations originating from the pituitary-adrenal feedback loop [3]. Its dynamics can be affected by stressors, exogenous CORT, and the estrous cycle. The phase *φ*_*C*_ is given by:

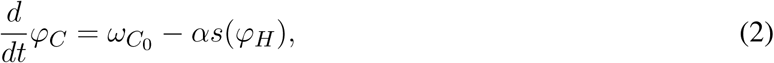

where 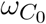 is the natural frequency of CORT ultradian oscillations, *s*(*φ*_*H*_) is a function accounting for a transient acute stressor (equal to zero in the absence of stress), and *α* is a scaling factor accounting for how strongly such stressor temporarily disrupts CORT ultradian rhythmicity. The amplitude *A*_*C*_ is given by:

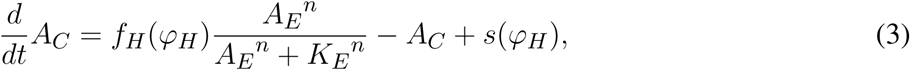

where *f*_*H*_(*φ*_*H*_) is a function representing hypothalamic circadian modulation. *A*_*E*_ is the amplitude of the estrous cycle (representative of the level of sex steroids) which modulates *A*_*C*_ through a Hill type function with coefficient *n* and half-maximum constant *K*_*E*_.

#### Pulse generator

Accounts for the activity of the GnRH pulse generator. Its frequency is modulated by stressors, CORT levels, and the activity of the KNDy network [4], which is in turn influenced by the phase of the estrous cycle. The phase *φ*_*PG*_ is given by:

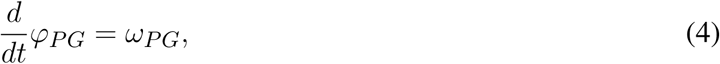

where *ω*_*PG*_ denotes the varying frequency of the pulse generator. This is given by:

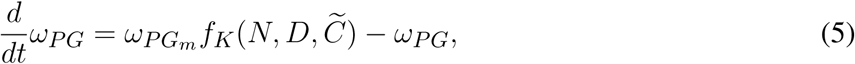

where 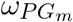 is the maximum frequency of the pulse generator and 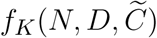 is a function accounting for the regulation from the KNDy network and CORT. Equations for the excitatory (*N*; e.g., Neurokinin B and glutamate) and inhibitory (*D*; e.g., Dynorphin) signals regulating the frequency of the KNDy network, and the slow genomic CORT effects 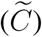 are given in the Supplementary Material.

#### Estrous cycle

Accounts for the activity of the reproductive cycle. The phase *φ*_*E*_ is given by:

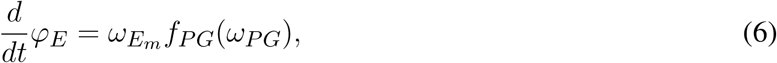

where *f*_*PG*_(*ω*_*PG*_) is a function accounting for the effects of the pulse generator and 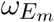 is the maximum frequency of the estrous cycle. The amplitude *A*_*E*_ is given by:

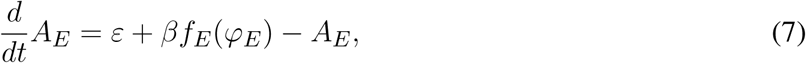

where *ε* is the basal activity of the estrous cycle, *f*_*E*_(*φ*_*E*_) is a function representing the effects of the estrous cycle, and *β* is a scaling factor accounting for the strength of such effects.

### 2.2 Computer simulations and parameter estimation

To simplify our analysis, CORT oscillations were normalised to the maximum levels observed in physiological conditions. That is, the CORT amplitude, which is modulated by the circadian drive, spans the range between 0 and 1 unless stressors or exogenous CORT act upon it. Similarly, the activity of the PG was represented by normalised oscillations, with a frequency that changes periodically according to the different stages of the estrous cycle. The model equations were numerically solved and analysed in MATLAB R2020a using ode45 routines. Details of the mathematical model development and parameter values are described in the Supplementary Material. The model parameters were estimated from the literature where available and manually calibrated to reproduce experimental observations of CORT and reproductive rhythms in rodents.

## 3 RESULTS

### 3.1 Normal physiological HPA and HPG rhythms

We calibrate the model parameters to reproduce physiological HPA and HPG rhythms observed in rats [1]. Accordingly, our model simulates CORT oscillations with a 75 min period, while the amplitude of these ultradian pulses is modulated in a circadian manner, reaching a maximum at the start of the dark period (Fig. 2A). Furthermore, one full estrous cycle lasts approximately 4 days, matching the average cycle length measured in rats [30]. A recent study using fibre photometry calcium imaging from arcuate kisspeptin neurons in mice revealed the dynamic modulation of GnRH pulse frequency along the estrous cycle [31]. Following these findings, the activity of the PG in the model remains inhibited (below 1 pulse/hr) during the post-ovulatory, estrous phase, rises steeply at the start of metestrus, and levels off at 2 pulses/hr for the rest of the cycle (Fig. 2B).

**Figure 2.**
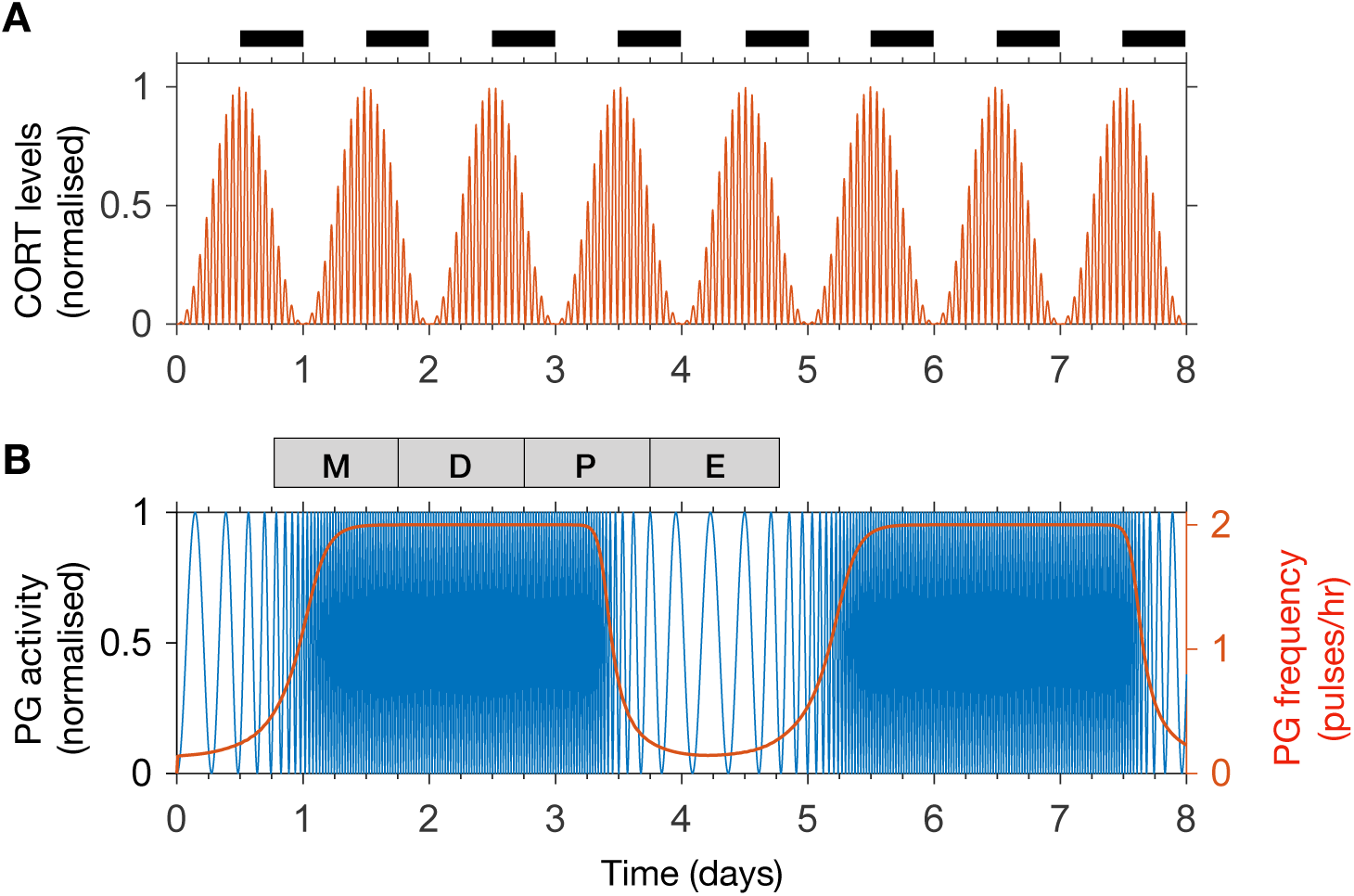
The model reproduces physiological rhythms in the HPA and HPG axis. **(A)** Normalised CORT levels as a function of time. The light-dark cycle is represented with intermittent black bars on the top. **(B)** Normalised pulse generator activity (blue) and pulse generator frequency (red) as a function of time. The phases of the estrous cycle are marked on the top: estrus (E); metestrus (M); diestrus (D); and proestrus (P).

### 3.2 Recovery of CORT dynamics following ovariectomy

Previous findings suggest that gonadal steroids are integral to the increased CORT levels seen in females compared to males. This has been demonstrated by showing the effects of oestrogen replacement in recovering physiological CORT levels following ovariectomy in rats [12]. We investigate the dynamic effects of these hormones by simulating the inhibition of HPA axis activity resulting from ovariectomy (OVX) and its restitution following 17*β*-oestradiol (E_2_) replacement. In the model, this is achieved by replacing the influx term in the right hand side of Eq. 7 by a constant term representing a drop in E_2_ levels following OVX (causing *A*_*E*_ to drop down to a constant level of 2% from the estrous peak) and by replacing the periodic sensitivity of the KNDy network to the estrous phase by a constant low value (Supplementary Material). The model predicts a drop in CORT levels down to ∼30% from its physiological value without loss of circadian or ultradian CORT rhythmicity while keeping the PG frequency at a high constant value of 2 pulses/hr (Fig. 3A). We then simulated the effects of an E_2_ pellet on OVX rats by increasing the constant value of the influx term in the right hand side of Eq. 7 (98% from physiological *A*_*E*_) and increasing the sensitivity of the KNDy network to the estrous phase (*φ*_*E*_) by a constant value (Supplementary Material). In agreement with [12], the model predicts recovery of physiological CORT levels without loss of circadian or ultradian CORT rhythmicity while marginally reducing the PG frequency just below 2 pulses/hr (Fig. 3B).

**Figure 3.**
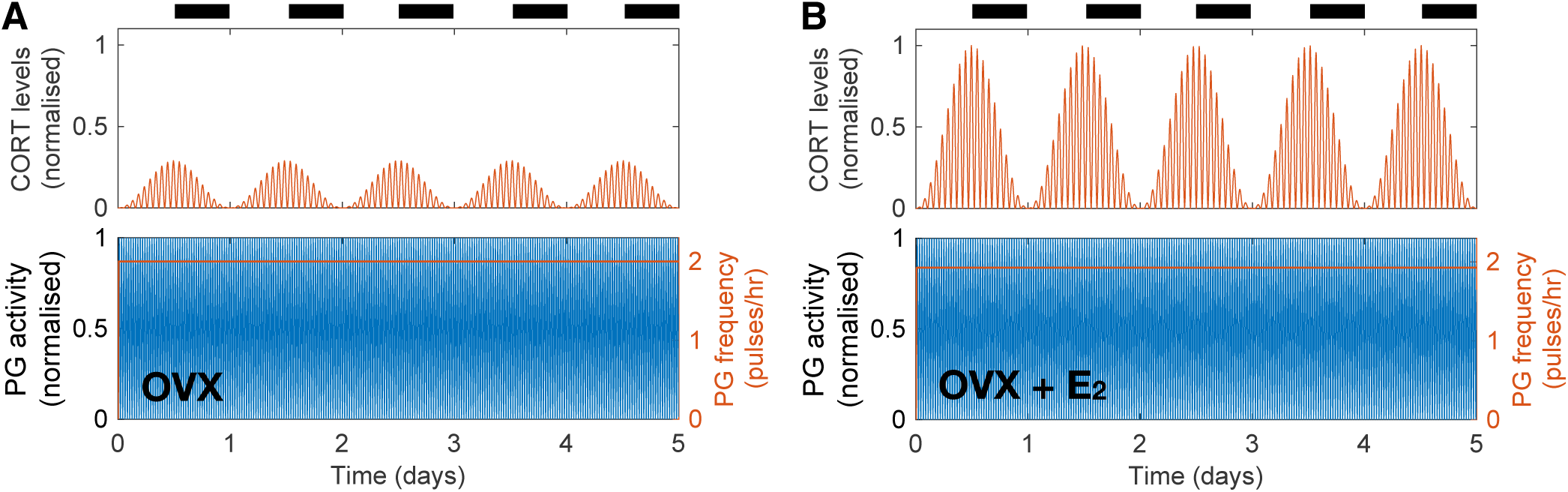
The model explains how E_2_ replacement recovers physiological CORT levels in OVX rats. **(A)** Simulated OVX reduced CORT oscillations down to ∼30% of the maximum physiological levels while keeping a constant high PG activity. **(B)** Simulated OVX+E_2_ recovered CORT oscillations to physiological levels while keeping a constant high PG activity.

### 3.3 Estradiol-mediated inhibition of HPG dynamics by high CORT doses

In a recent study, Kreisman and co-workers [8] investigated the effect of chronic CORT administration on LH pulsatility and demonstrated the importance of gonadal steroid hormones in mediating the inhibitory effect of CORT on the HPG axis. The study showed that a pellet delivering a high dose of CORT over 48 hrs in OVX mice has no effect on LH pulsatility, whereas a significant reduction of LH pulse frequency is observed in OVX animals treated with a 17*β*-estradiol silastic implant (OVX+E_2_). In our model, we accounted for the OVX and OVX+E_2_ scenarios as described in the previous section, while the constantly high CORT levels were achieved by replacing the effective CORT levels modulating the KNDy network by a constant high value estimated from [8] (see Supplementary Material).

Figure 4 illustrates the differential effect of chronically elevated CORT levels on the GnRH pulse generator frequency in OVX versus OVX+E_2_ animals. In the case of OVX animals, elevated CORT levels do not alter the frequency of the pulse generator, whereas in OVX animals treated with estradiol the frequency is halved for as long as CORT levels are elevated. This effect is linked to the modulation of the GnRH pulse generator by gonadal steroids, which sensitise the system to inhibitory signals such as CORT or acute stressors as we show below (Fig. 4B).

**Figure 4.**
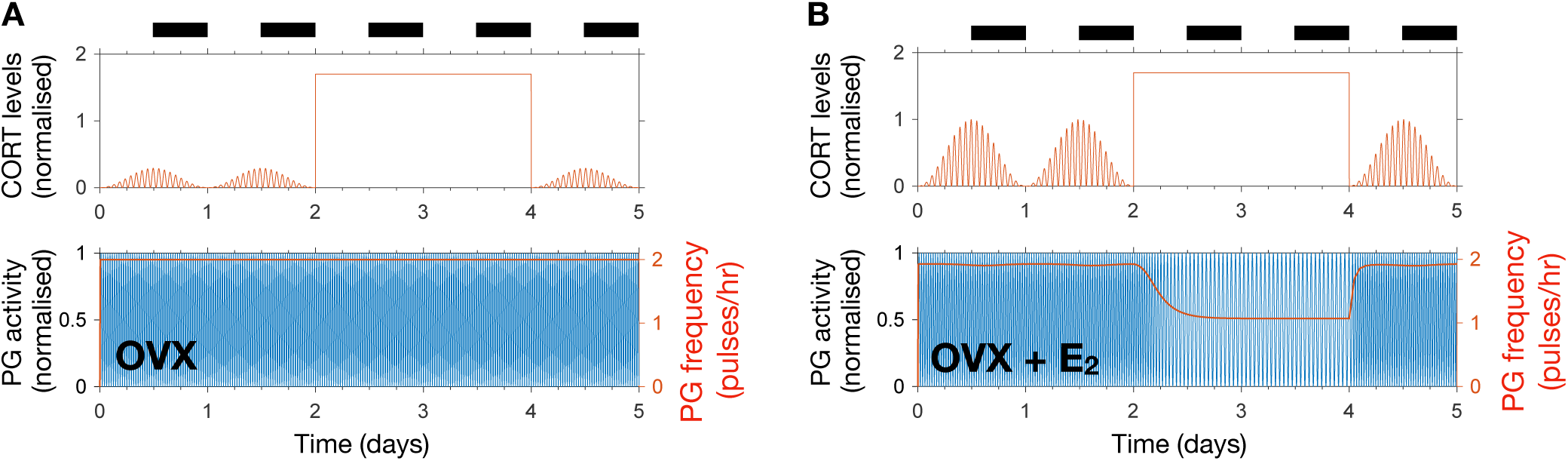
The model reproduces estradiol-mediated inhibition of PG activity following high doses of CORT. **(A)** High exogenous CORT over 48 hrs does not affect the PG dynamics in OVX mice. **(B)** In the presence of estradiol, high CORT doses temporarily reduce PG activity in OVX mice.

### 3.4 Acute stress effects on the HPA and HPG axes depend on the estrous cycle phase

To study the effect of acute stress on the dynamics of the HPA and HPG axes, we extend the model to include transient stress-related neuronal inputs affecting both axes [11]. In our model, we account for these transient inputs by simulating a 2 hr square pulse of amplitude 1, equivalent to a restraint stressor causing a CORT increase from its circadian nadir up to its circadian peak [32]. The stressor affects the phase and amplitude of the CORT rhythm (function *s*(*φ*_*H*_) in Eqs. 2 and 3) as well as the frequency of the GnRH pulse generator 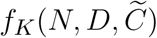 in Eq. 5 and Supplementary Material).

Figures 5A and 5B illustrate the effect that 2 hr of stress activation has on the dynamics of the HPA and HPG axes when applied at different times along the cycle. Both CORT and GnRH pulse generator responses are dependent on the timing of the input pulse (Fig. 5C). The amplitude of the CORT response shows a circadian dependency with stressors delivered during the circadian peak eliciting a stronger response. The GnRH pulse generator frequency response to acute stressors depends on the phase of the estrous cycle. In particular, the frequency of the pulse generator appears most sensitive to stressors during estrus to early diestrus phases, with little or no effect during the mid-cycle phase. This differential effect of acute stress on the frequency of the GnRH pulse generator activity highlights the cycle dependent modulation of the pulse generator dynamics, which makes the pulse generator more robust to perturbations in the diestrus and proestrus phases (Fig. 5D).

**Figure 5.**
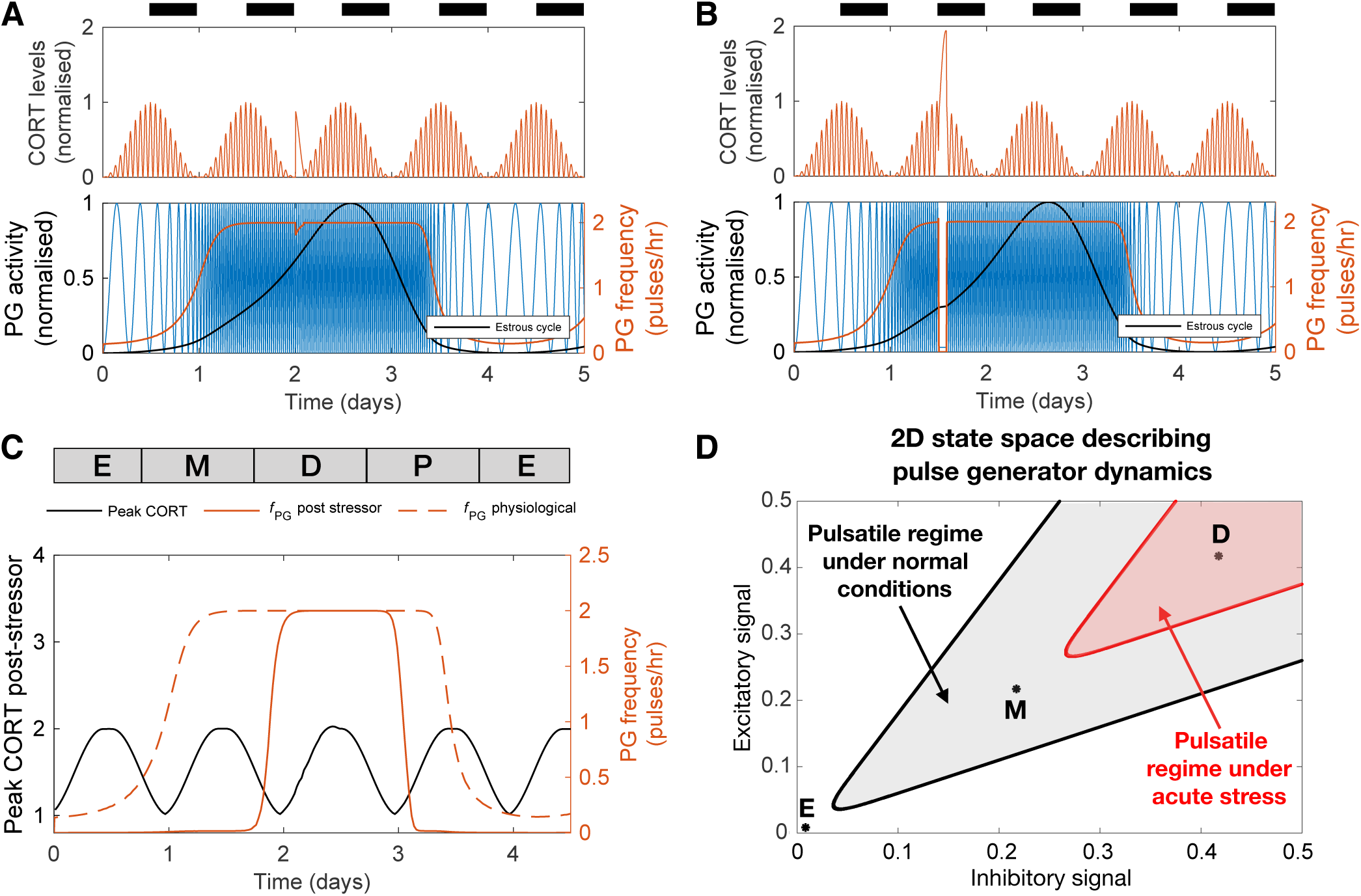
The effect of acute stress on the dynamics of the HPA and HPG axes. **(A-B)** CORT levels and PG activity in response to a transient (2 hr long) stressor initiated at two different times. **(C)** Peak CORT levels (black line) and mean PG frequency (continuous red line) elicited by a 2 hr long acute stressor as a function of the time at which the stressor arrives during the estrous cycle. The PG frequency without any stress perturbation is shown for comparison (dashed red line). **(D)** State space diagram describing the effect of acute stress on the dynamics of the pulse generator. Points mark different stages along the estrus cycle: estrus midpoint (E); metestrus midpoint (M); and diestrus midpoint (D). The shaded gray area denotes the region of the state space corresponding to frequencies above 1 pulse/hr under normal physiological conditions. Acute stress shrinks this region (red shaded area), but the dynamics of the pulse generator maintains robustness to perturbations during the diestrus phase.

### 3.5 CORT excess increases the length of the estrous cycle and modulates responses to acute stressors

Last, we used the model to predict the effects of high excess CORT (hiCORT) —mimicking levels expected to be observed in people with hypercortisolism— on the estrous cycle. To do this, we considered the increase in baseline and maximum CORT amplitude with respect to physiological levels in humans [33] and implemented the equivalent increase ratios for our simulations of CORT dynamics in rodents (Supplementary Material). Evidence from high frequency sampling in humans shows hypercortisolism is associated with a reduction in the ultradian period of CORT oscillations [34]. Accordingly, we also adjusted this parameter when modelling hiCORT, while keeping circadian oscillations and all other parameters unchanged. Our simulations predict an increase in the period of the estrous cycle from a physiological value of T_*phys*_ = 4.2 days up to T_*hiC*_ = 5.1 days under hiCORT, which is equivalent to a ∼21% increase in the estrous cycle length (Fig. 6A and C).

**Figure 6.**
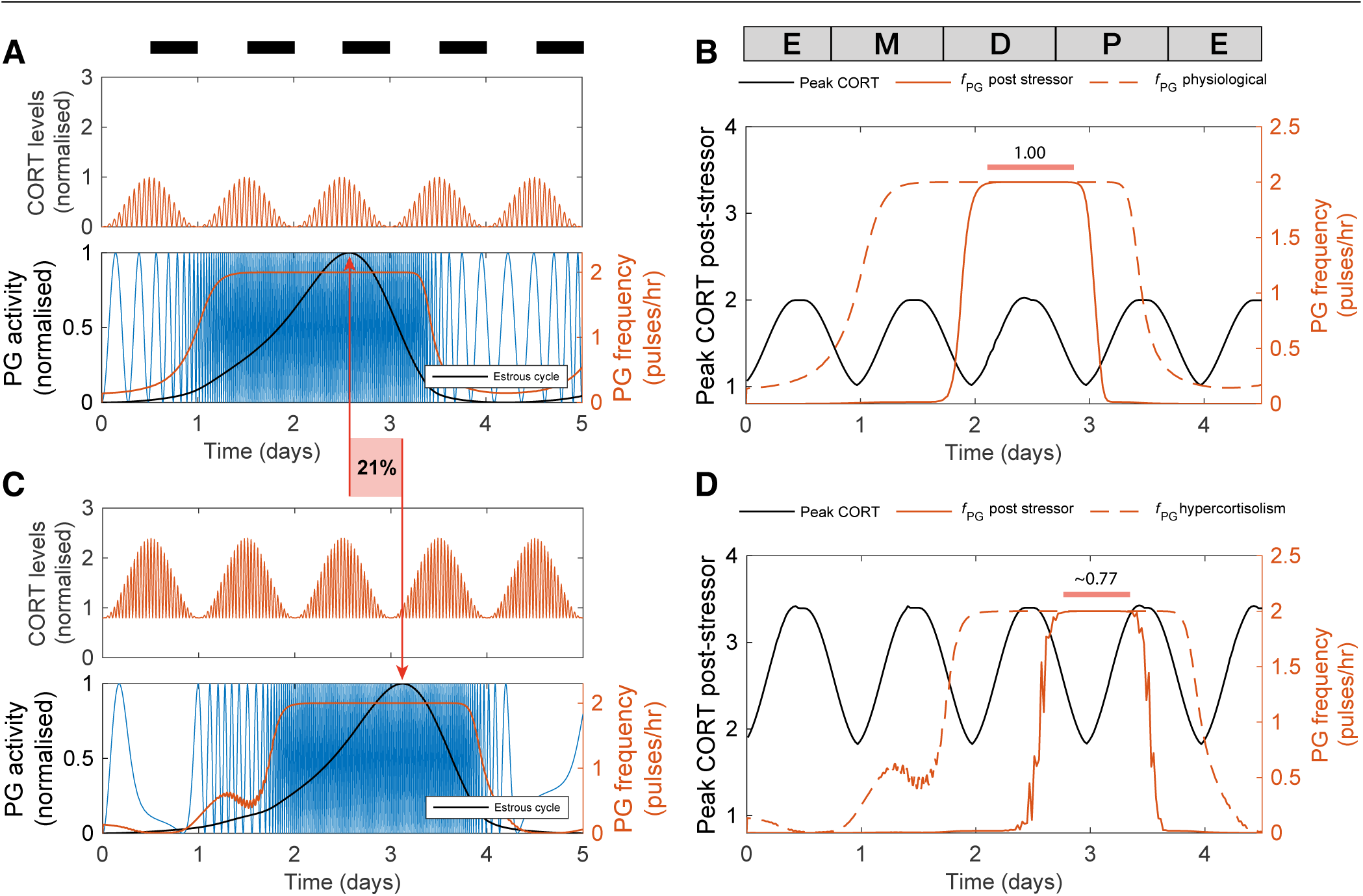
Enduring and transient dynamic changes under hiCORT. **(A)** CORT and PG rhythms in physiological conditions. **(B)** Mean PG frequency and maximum CORT levels elicited by 2 hr long hypothalamic stressors arriving at different times across the estrous cycle. **(C)** Under hiCORT, the range of CORT levels is increased and the estrous cycle peak is delayed by ∼ 21% compared to physiological values. **(D)** Mean PG frequency and maximum CORT elicited by 2 hr long stressors under hiCORT. In this scenario, the period in which PG activity remains unchanged by stressors starts about half a day later and is shortened compared to normal physiological conditions.

We then used the model to investigate the transient changes in the GnRH pulse generator frequency and CORT amplitude elicited by exogenous acute stressors under physiological conditions and hiCORT. In particular, we look at the effects of the timing of stressors within the estrous cycle. To do this, we calculated frequency and amplitude response curves by simulating a 2 hr long stressor elicited at different stages across the estrous cycle using 30 min time steps. In the physiological scenario (Fig. 6B), the model predicts that acute stressors suppress PG activity during most of the estrous cycle except during the diestrus and early proestrus phases. These stressors also elicit an increase in peak CORT levels to a range between 1 and 2. While the model under the hiCORT scenario predicts a similar behaviour, the region where PG activity remains unaffected by acute stressors is reduced and delayed by about half a day compared to its physiological counterpart (Fig. 6D and Fig. S2). This is not surprising considering that the model also predicts that hiCORT prolongs the estrous cycle. Regarding the CORT response to stressors under hiCORT, our model predicts a ∼2 to ∼3.5 increase in CORT levels compared to the normal physiological scenario. This is due to a compounded effect of CORT surges over an excess CORT baseline.

## 4 DISCUSSION

We developed and studied a mathematical model that integrates components of the stress and reproductive axes at different spatial and temporal scales, from the molecular intricacies of the KNDy network, to GnRH and CORT oscillations, up to the estrous cycle (Fig. 1A). Previous mathematical models of the HPA and HPG axes either focus on a specific process within an axis, or consider them as a whole, but isolated from each other [6, 2, 3, 4]. In contrast, our model integrates these neuroendocrine axes by considering the complex interactions between them as a network of interlocked oscillators, hence enabling us to integrate different physiological observations and experiments into a single coherent theoretical framework and study the effect of transient perturbations on the overall dynamics. In particular, our model postulated a network architecture (Fig. 1B) that reflects physiological observations of ultradian and circadian CORT rhythms, as well as ultradian and infradian rhythms of the GnRH pulse generator (Fig. 2). The model reproduced the effects of ovarian hormone removal (OVX) and restitution (E_2_) on the HPA and HPG axes dynamics, both under physiological conditions [12] (Fig. 3) and under exogenous CORT excess [8] (Fig. 4).

In addition to these slow timescale perturbations, we also investigated the fast timescale perturbations elicited by acute stressors. Our model predicted that exogenous stress perturbations not only cause transient increases in CORT levels, but also transiently inhibit GnRH pulse generator activity with the magnitude of this inhibition being dependent on the estrous and circadian phases (Fig. 5). This has important implications about understanding how the timing of a stressor affects its ability to temporarily suppress the GnRH/LH ovulatory surge. According to our model, the pulse generator activity is robust to stress perturbations arriving between the diestrus and early proestrus stage, but is fragile to stressors arriving at estrus and metestrus stages (Fig. 5C). Uncovering the origin of this robustness is beyond the scope of our phenomenological model, but we can speculate that molecular mechanisms ensure the resilience of the reproductive cycle during the key stages leading to ovulation. While our model suggests that the GnRH/LH surge should be delayed under frequent exposure to stressors, if the exposure occurs too close to the proestrus stage then these resilience molecular mechanisms ensure the surge continues as normal and triggers ovulation [35, 36].

We also used the model to investigate the potential detrimental effects on fertility elicited by chronic hypercortisolism. Our model predicted that hiCORT delays the increase in activity of the GnRH pulse generator, effectively prolonging the estrous cycle (Fig. 6A and C). While evidence suggests that HPA axis hyperactivity—and specifically, increased circulating glucocorticoids—are unlikely to be the sole mechanism behind stress-induced reproductive dysfunction [19], our simulations show the cycle length depends on the GnRH pulse generator’s sensitivity to CORT (Fig. S1A). Thus, our model provides insight into how for example a hyper-sensitized HPG axis may explain amenorrhea secondary to high serum cortisol levels [16, 17, 18]. Interestingly, our model predicted that a period of robustness of the GnRH pulse generator in the presence stressors is preserved under hiCORT, albeit the robust period occurs about half a day later in the cycle and is shorter in duration. Our model simulations of pulse generator activity suggest that prolonging the estrous cycle as predicted under hiCORT arises from a combination of longer estrus and metestrus stages while diestrus and proestrus stages are shortened (Fig. 6B and D).

Our model considers essential features of HPA and HPG axes oscillators in a phenomenological way. This approach facilitates the simulation of a range of physio-pathological scenarios, but inevitably imposes certain limitations. In contrast to mechanistic models where parameters are often linked to chemical kinetic rates, the parameters in our model represent natural and maximum frequencies, phase relationships, as well as the coupling strengths between oscillators and sensitivities to perturbations. While our phenomenological approach limits the ability of the model to support discovery of specific molecular mechanisms, it can be used to suggest experiments that explore systems level properties involving both neuroendocrine axes. For example, evidence shows that in addition to exhibiting circadian and ultradian fluctuations, CORT levels also change across the estrous cycle, with maximum levels around the diestrus and proestrus phase [37, 38, 39]. While our model lacks the level of detail to describe the molecular mechanisms that underpin estrous changes on CORT, it does suggest this is mediated by a regulatory link from the estrous oscillator to the CORT oscillator, thus inferring that ovarian steroids may be the culprit of estrous regulation of CORT instead of the hypothalamic GnRH pulse generator (Fig. 1). In our model, we only explored the scenario where the strength of this regulatory link allows for strong perturbations (e.g., stressors, OVX, E_2_) in the estrous oscillator to have an impact on the CORT dynamics, but not from milder estrous regulation of CORT levels (Fig. S1). We speculate that combining mechanistic modelling with experimental physiology to investigate the effects of estradiol and progesterone on CORT may uncover the origins of its estrous cycle modulation. The experiments could test the dosing effect, timing, and combined sensitivity of gonadal steroids on circadian CORT levels across the estrous cycle. The mechanistic model could in turn help understand the robustness of such regulatory mechanism to perturbations [35], and predict the scenarios in which chronodisruptions would lead to disease.

We believe that the first generation mathematical model presented here could be used to inform further investigations into the timing of stress perturbations in reproductive health, including dysregulations induced by strenuous exercise [16], mood disorders [40], as well as clinical interventions such as *in vitro* fertilisation [41]. We anticipate that healthcare technologies such as wearable devices and smartphone apps collecting vast amounts of data on body rhythms, together with computer algorithms characterising inter-individual variability, will help refine and personalise neuroendocrinological models [42, 43].

## AUTHOR CONTRIBUTIONS

E.Z., M.V., X.F.L., J.R.T., S.L.L., K.O., and K.T.A. conceived and designed the study. E.Z., M.V., T.Z., J.T., J.J.W., and K.T.A. developed the mathematical model. E.Z., M.V. and T.Z. performed the analytical calculations and computer simulations. E.Z., M.V. and T.Z. wrote the manuscript with support of all authors.

## SUPPLEMENTAL DATA

See annex Supplementary Material for auxiliary equations and coupling functions, tables of model parameter values, and supplementary figures.

## DATA AVAILABILITY STATEMENT

The computer code generated through this study is the subject of current research and is available upon request.

