## Supplementary Material for "Dynamic hormone control of stress and fertility"

The state variables of our model are listed in Table S1. The main model equations for all state variables can be found in the manuscript. In this supplement we provide the auxiliary equations and functions used to represent the external inputs and the coupling between model variables. We also include tables of model parameter values for all scenarios simulated and supplementary figures.

| Variable | Description |
|---|---|
| | State Variables |
| $\varphi_H$ | Phase of hypothalamic oscillator. |
| $\varphi_C$ | Phase of CORT oscillator. |
| $\varphi_E$ | Phase of estrous cycle. |
| $\varphi_{PG}$ | Phase of PG oscillator. |
| $\omega_{PG}$ | Frequency of PG oscillator. |
| $A_C$ | Amplitude of CORT oscillator. |
| $A_E$ | Amplitude of estrous cycle. |
| | Auxiliary Variables |
| $N$ | Activity of excitatory signals within the KNDy network (e.g., Neurokinin-B, glutamate). |
| $D$ | Activity of inhibitory KNDy signals within the KNDy network (e.g., Dynorphin). |
| $\widetilde{C}$ | Slow CORT activity due to genomic effects. |

**Table S1.** Model state variables.

### 1 AUXILIARY EQUATIONS AND COUPLING FUNCTIONS

#### 1.1 Acute stressor

An acute stressor $s(\varphi_H)$ is modelled as a squared pulse defined by the product of two discrete Heaviside step functions:

$$s(\varphi_H) = p_A H(\varphi_H - \varphi_s) H(\varphi_s + \theta_s - \varphi_H), \tag{S1}$$

where $p_A$ is the pulse amplitude, $\varphi_H$ is the circadian phase, $\varphi_s$ is the phase at which the pulse starts, and $\theta_s$ is the duration of the pulse. To simulate normal physiological rhythmicity with no pulse stressor, we simply set $p_A = 0$. An acute stressor $s(\varphi_H)$ affects CORT phase $\varphi_C$, amplitude $A_C$, and the pulse generator frequency $\omega_{PG}$ through its coupling to the KNDy network and CORT (see Eqs. S9 and S10).

#### 1.2 Hypothalamic circadian regulation of CORT

The hypothalamic circadian regulation of the CORT amplitude $A_C$ is modeled by:

$$f_H(\varphi_H) = a + b \sin^2(\varphi_H), \tag{S2}$$

where $a$ and $b$ represent the CORT baseline and the maximum range over the baseline, respectively. These control parameters can be used to switch between simulations of normal physiological CORT dynamics ($a = 0$, $b = 1$) and hypercortisolism ($a = 0.8$, $b = 1.6$) (Vagnucci, 1979). The ultradian period under hypercortisolism has been adjusted following recent high-frequency sampling studies (Van Aken et al., 2005). For physiological CORT levels we set $\omega_{C_0} = \pi/75$, which corresponds to an ultradian period of ≈19 min. For hypercortisolism we set $\omega_{C_0} = \pi/125$, which corresponds to an ultradian period of ≈12 min (Tables S2 and S3).

#### 1.3 Estrous cycle regulation

We model the estrous cycle regulation of its amplitude and its effects on the KNDy network through a skewed sinusoidal function of the phase $\varphi_E$. This is given by:

$$f_E(\varphi_E) = \begin{cases} \sin^2\left(\varphi_E - \sigma \sin^2(\varphi_E)\right), & \text{Normal physiological} \\ 0.99, & \text{OVX} \\ 0.2, & \text{OVX+E}_2 \end{cases} \tag{S3}$$

where $\sigma$ is the skewness of the estrous cycle. We fix $f_E(\varphi_E)$ to a constant value to simulate the OVX and OVX+$E_2$ scenarios (Kreisman et al., 2019; Seale et al., 2004). Note that in those cases, parameters $\varepsilon$ and $\beta$

in the equation for $A_E$ also need to change as indicated in Table S3 to reflect estrous activity expected in the diestrus and proestrus phase.

##### 1.4 Pulse generator regulation of the estrous cycle

The pulse generator frequency $\omega_{PG}$ modulates the progression through the estrous cycle by acting on the estrous phase $\varphi_E$ as:

$$f_{PG}(\omega_{PG}) = e^{-\lambda}, \quad \text{with} \quad \lambda = \frac{\left(\omega_{PG} - \omega_{PG}^{\mu}\right)^2}{2{\omega_{PG}^{\sigma}}^2}, \tag{S4}$$

where $\omega_{PG}^{\mu}$ and $\omega_{PG}^{\sigma}$ are the mean and spread of a bell-shaped response of the estrous phase to the PG frequency. Such a model accounts for cycle disruptions when the pulse generator frequency is set either too low or too high (Knobil et al., 1980; Pohl et al., 1983).

##### 1.5 KNDy network and CORT regulation of the pulse generator

The frequency of the pulse generator $\omega_{PG}$ is governed by the activity of the KNDy network and the concentration of CORT. To account for this, we model excitatory ($N$; e.g., Neurokinin-B, glutamate) and inhibitory signals ($D$; e.g., Dynorphin) within the KNDy network (Voliotis et al., 2019) as:

$$\frac{d}{dt}N = f_E(\varphi_E) - N, \tag{S5}$$

$$\frac{d}{dt}D = f_E(\varphi_E) - D, \tag{S6}$$

where $f_E(\varphi_E)$ accounts for the periodic modulation of the KNDy network activity occurring through the progression of the estrous cycle (see S3).

The CORT concentration $C$ is a function of the state variables $\varphi_C$ (CORT phase) and $A_C$ (CORT amplitude), and is given by:

$$C = A_C \sin^2(\varphi_C), \tag{S7}$$

where the ultradian rhythm of CORT is represented by $\sin^2(\varphi_C)$ and the circadian rhythm is contained in $A_C$. The CORT concentration $C$ thus fluctuates with circadian and ultradian rhythmicity.

We expect CORT to exert its effect on the pulse generator (PG) at the genomic level (e.g., regulating the expression of components in the KNDy network) (Ayrout et al., 2019). Since genomic effects are typically slower than systems-level CORT effects considered in the model, we introduce a new CORT variable $\widetilde{C}$ to account for for slow genomic effects. Thus, $\widetilde{C}$ filters out the fast dynamics of $C$ and is described by:

$$\frac{d}{dt}\widetilde{C} = \delta(C - \widetilde{C}), \tag{S8}$$

where $\delta$ defines the slow timescale. In the scenarios where we simulate exogenous CORT levels, $C$ in Eq. S8 is replaced by a constant $p_C$ equivalent to the dose delivered by a pellet (Table S3).

Considering the activity of $N$, $D$ and the filtered CORT $\widetilde{C}$, we can model their effect on the pulse generator frequency $\omega_{PG}$ as (Voliotis et al., 2019):

$$f_K(N, D, \widetilde{C}) = \frac{e^{h\left(x-g(s,\widetilde{C})\right)}}{1+e^{h\left(x-g(s,\widetilde{C})\right)}}\frac{e^{h\left(y-g(s,\widetilde{C})\right)}}{1+e^{h\left(y-g(s,\widetilde{C})\right)}}, \quad \text{with} \quad x = D - \frac{N}{2} \quad \text{and} \quad y = N - \frac{D}{2}, \tag{S9}$$

where $h$ is a parameter that accounts for the overall contribution of $N$ and $D$ to the PG frequency. The function $g(s, \widetilde{C})$ embedded in Eq. S9 takes the form:

$$g(s, \widetilde{C}) = \frac{1}{h}\left(1 + m\frac{s(\varphi_H)^n}{{K_s}^n + s(\varphi_H)^n} + l\frac{\widetilde{C}^n}{{K_C}^n + \widetilde{C}^n}\right). \tag{S10}$$

where $K_s$ accounts for the PG frequency sensitivity to an acute stressor $s(\varphi_H)$ (Eq. S1), $K_C$ accounts for the PG frequency sensitivity to filtered CORT levels, and $m$ and $l$ account for the overall contribution of acute stressors (see S1) and the filtered CORT signal to the PG frequency, respectively.

#### 2 MODEL PARAMETERS

The oscillator frequencies, sensitivities and other parameter values were fixed manually to reproduce observations in the normal physiological scenario and the experiments by Seale et al. (2004) and Kreisman et al. (2019). The parameter values for physiological and phyiso-pathological scenarios are listed in Tables S2 and Table S3.

| Parameter | Description | Value |
|---|---|---|
| $\omega_{H_0}$ | Circadian natural frequency ($T = 24$ h). | $\pi/(24*60)$ |
| $\omega_{C_0}$ | Ultradian CORT natural frequency ($T = 75$ min). | $\pi/75$ |
| $\omega_{E_m}$ | Estrous cycle maximum frequency ($T_{\text{min}} = 3$ days). | $\pi/(24*60*3)$ |
| $\omega_{PG_m}$ | Pulse generator maximum frequency ($T_{\text{min}} = 30$ min). | $\pi/30$ |
| $\omega_{PG}^{\mu}$ | Mean of bell-shaped response of the Estrous cycle to PG frequency. | $\pi/30$ |
| $\omega_{PG}^{\sigma}$ | Spread of bell-shaped response of the Estrous cycle to PG frequency. | $\pi/60$ |
| $\alpha$ | Strength of stressor disruption on $\varphi_C$ dynamics. | 0.05 |
| $\beta$ | Strength of Estrous cycle phase on its amplitude. | 0.9 |
| $\delta$ | Timescale reduction of filtered CORT dynamics. | 0.005 |
| $\varepsilon$ | Basal activity of the Estrous cycle. | 0.1 |
| $\sigma$ | Estrous cycle skewness. | 0.5 |
| $\phi_s$ | Starting phase of the pulse stressor. | $2\pi$ |
| $\theta_s$ | Phase duration of the pulse stressor. | $\pi/12$ |
| $K_E$ | CORT amplitude sensitivity to the Estrous cycle amplitude. | 0.025 |
| $K_s$ | PG frequency sensitivity to pulse stressor. | 0.6 |
| $K_C$ | PG frequency sensitivity to $\widetilde{C}$. | 1.7 |
| $E_{lo}$ | Estradiol levels during the diestrous phase of the cycle. | 0.02 |
| $E_{hi}$ | Estradiol levels during the proestrous phase of the cycle. | 0.98 |
| $p_A$ | Acute stress pulse amplitude. | 1 |
| $h$ | Overall contribution of KNDy network to PG frequency. | 50 |
| $m$ | Contribution of acute stressor to PG frequency. | 13 |
| $l$ | Contribution of CORT levels to PG frequency. | 6 |
| $n$ | Hill coefficient. | 4 |

**Table S2.** Parameter values for the physiological scenario.

| Parameter | Stressor | OVX | OVX+$E_2$ | CORT pellet | Hypercortisolism |
|---|---|---|---|---|---|
| $\omega_{C_0}$ | $\pi/75$ | $\pi/75$ | $\pi/75$ | $\pi/75$ | $\pi/125$ |
| $a$ | 0.8 | 0 | 0 | 0 | 0 |
| $b$ | 1.6 | 1 | 1 | 1 | 1 |
| $\beta$ | 0.9 | 0 | 0 | 0.9 | 0.9 |
| $\varepsilon$ | 0.1 | $E_{lo}$ | $E_{hi}$ | 0.1 | 0.1 |
| $p_C$ | - | - | - | 1.7 | - |

**Table S3.** Parameter values for physio- pathological simulations.

#### 3 SUPPLEMENTARY FIGURES

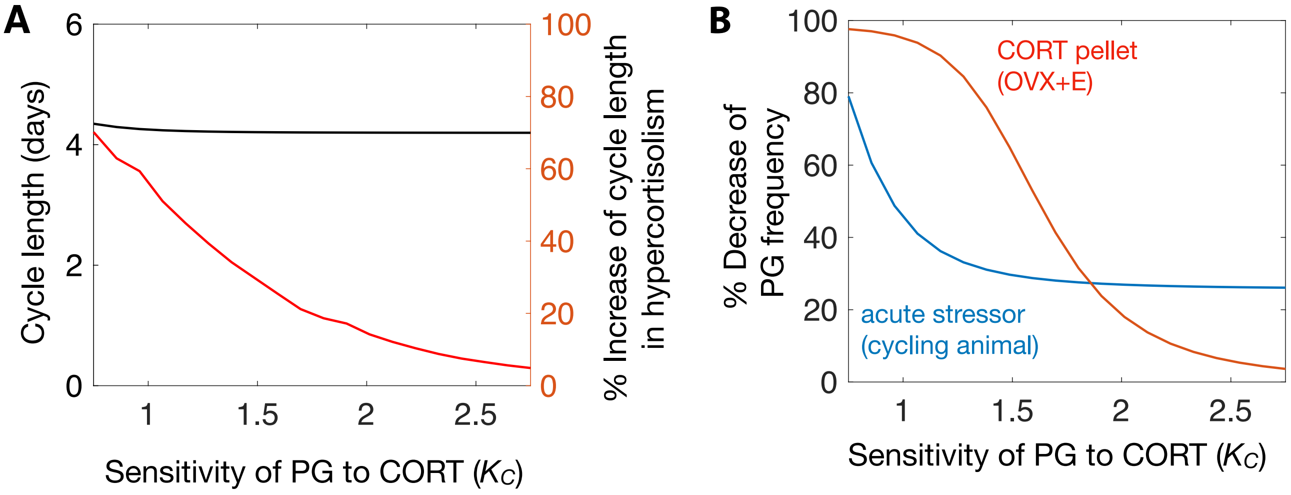


**Figure S1.** Sensitivity of GnRH pulse generator activity to CORT. **(A)** Estrous cycle length under normal physiological CORT levels and percentage increase of cycle length under hypercortisolism as a function of $K_C$. **(B)** Percentage decrease of PG frequency compared to the normal physiological scenario elicited by an acute stressor and by a CORT pellet as a function of $K_C$.

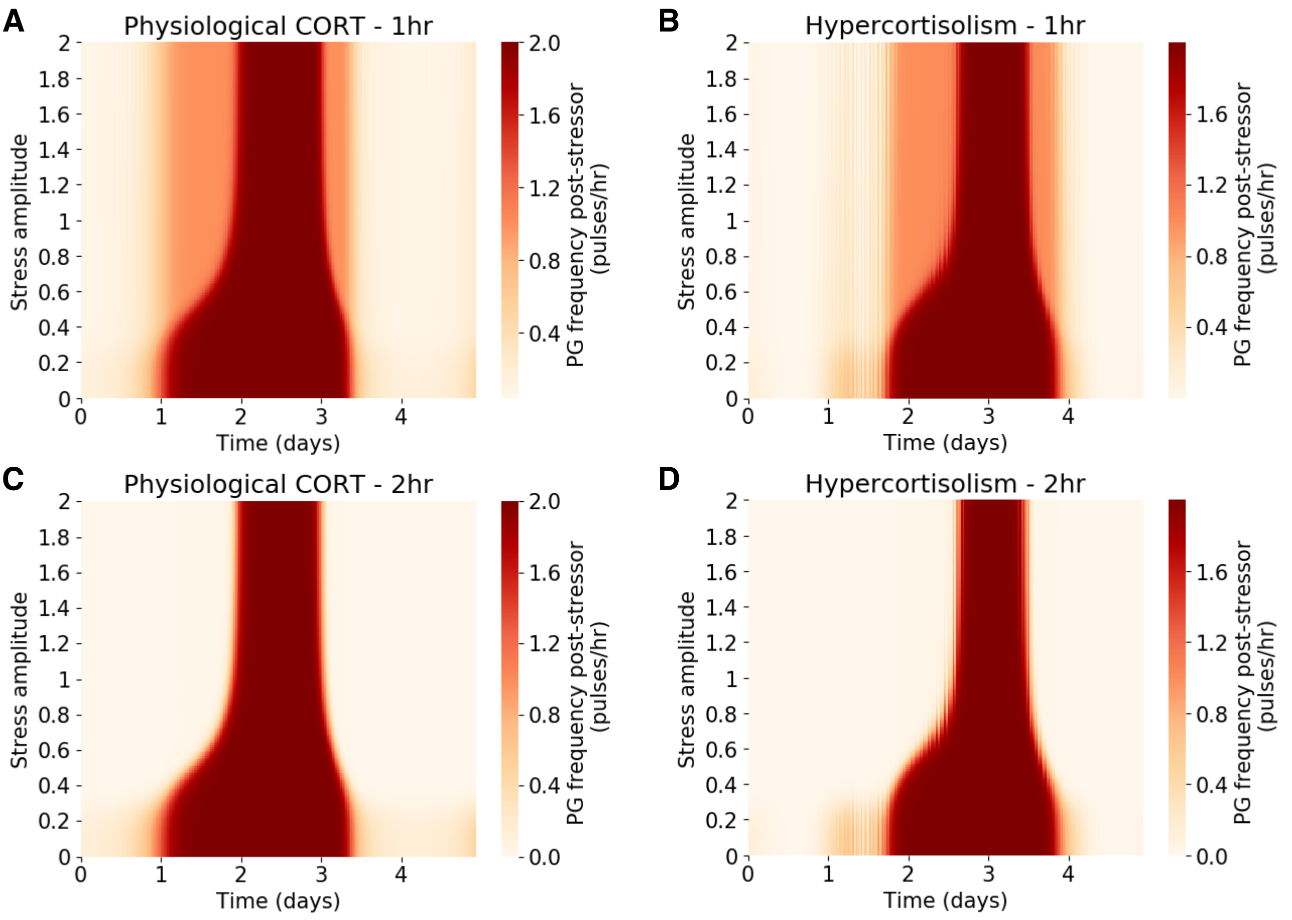


**Figure S2.** Average pulse generator frequency during an acute stressor as a function of stress amplitude $p_a$ and timing of stressor onset $\Phi_s$ (here shown in days). **(A)** PG frequency under normal physiological CORT levels during 1 hr long stressors. **(B)** PG frequency under hypercortisolism during 1 hr long stressors. **(C)** PG frequency under normal physiological CORT levels during 2 hr long stressors. **(D)** PG frequency under hypercortisolism during 2 hr long stressors. A stress amplitude of zero ($p_a = 0$) at the bottom of each panel corresponds to the normal physiological PG frequency without stress.

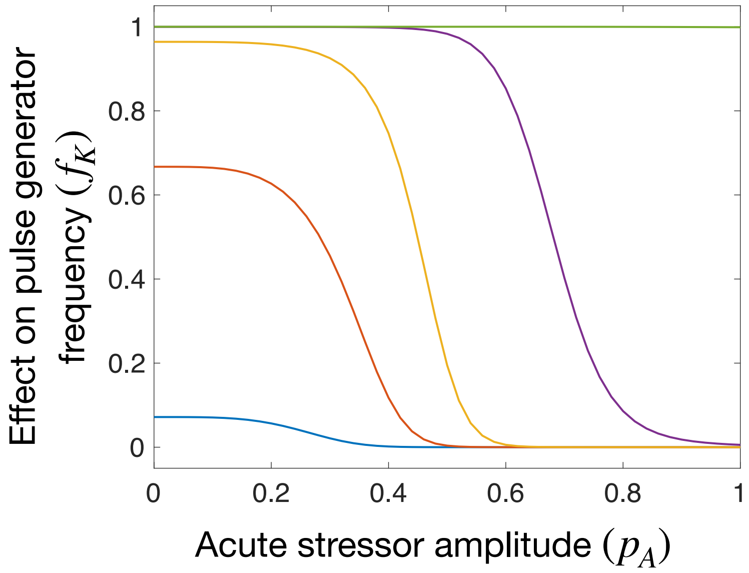


**Figure S3.** Effect of acute stressor amplitude ($p_A$) on the regulation of pulse generator frequency ($f_K$; see Eq. S9). Colored lines correspond to different levels of excitatory ($N$) and inhibitory signals ($D$) within the KNDy network: 0 (blue; corresponding to Estrus), 0.1 (red), 0.2 (yellow), 0.4 (purple; corresponding to Metestrus), 0.8 (green; corresponding to Diestrus). $\widetilde{C}$ = 0.3.
